# Morphology of softening and translucency in the stems of tomato and eggplant seedlings due to infection by *Ralstonia pseudosolanacearum* F1C1

**DOI:** 10.1101/2024.09.10.612199

**Authors:** Shuvam Bhuyan, Lakhyajit Boruah, Monika Jain, Shuhada Begum, Shubhra Jyoti Giri, Lukapriya Dutta, Sumi Kalita, Swarnava Chatterjee, Manabendra Mandal, Suvendra Kumar Ray

**Affiliations:** Department of Molecular Biology & Biotechnology, Tezpur University, Tezpur–784028, Assam, India

**Keywords:** *Ralstonia pseudosolanacearum*, Tomato, Eggplant, Symptoms, Stem morphology

## Abstract

*Ralstonia pseudosolanacearum* F1C1 is a soil-borne phytopathogenic bacterium with a broad host range that infects several economically important crops. This study primarily focuses on the infection of this phytopathogen in two such important crop seedlings: tomato and eggplant. The observations of the study reveal a complex set of symptoms that include drooping and blackening of the seedling stem, as well as blackening, chlorosis, and curling of cotyledon leaves. Notably, the symptom of stem softening and translucency is seen primarily in the water-submerged stem regions of both root- and leaf-inoculated seedlings. While the wild-type *R. pseudosolanacearum* F1C1 strain and its weakly virulent mutant *phcA*::Ω exhibited this phenotype in both tomato and eggplant seedlings, the virulence-deficient *hrpB*::Ω mutant did so only in a few eggplant seedlings but not in tomato seedlings. By investigating these unique pathological phenotypes in the infected seedlings, this study uncovers a unique symptom previously not reported in seedling inoculation experiments. The work also touches upon the distinct escape mechanisms exhibited by seedlings, revealing how some of its host plants might resist wilting despite infection, offering new avenues for future research. These findings contribute to the understanding of *R. pseudosolanacearum* pathogenesis, shedding light on its virulence and host response mechanisms.

## 1. Introduction

*R. pseudosolanacearum* causes a systemic infection by invading and spreading through the xylem vessels of its host plants (Vasse et al. 1995; Vasse et al. 2000; Digonnet et al. 2012). Considering the systemic nature of its infection, the disease symptoms are complex. In tomato (*Solanum lycopersicum*), wilting initially manifests in the younger, apical leaves at the top of a grown-up plant and gradually progresses downward to the older branches (Din et al. 2016). In contrast, in eggplant (*S. melongena*), wilting follows a bottom-up pattern, beginning with the older leaves and advancing toward the younger leaves (Gousset et al. 2005). In both hosts, this process eventually leads to the entire plant wilting and collapsing to the ground. While wilting is the most prominent symptom, other manifestations such as chlorosis and stunted growth are among the several disease symptoms that can be observed in its hosts (García et al. 2019). Apart from the well-documented disease phenotypes in grown-up plants observed through soil drenching or stem inoculation methods, we have recently demonstrated *R. pseudosolanacearum* F1C1 pathogenicity in cotyledon stage seedlings of eggplant and tomato using both cotyledon leaf clip and root inoculation methods (Singh et al. 2018; Kumar et al. 2017; Phukan et al. 2019). The pathogenicity in seedlings and the wilting phenotype are similar to grown-up plants, considering the virulence factors reported for grown-up plant infection have also been demonstrated to be required for the seedling’s infection (Singh et al. 2018; Kumar et al. 2017; Phukan et al. 2019).

The seedling model allows for close monitoring of both the progression of the infection and the disease, providing valuable insights into the pathogen’s behavior and effects. As the genome of the phytopathogen harbors numerous potential virulence functions with mechanisms yet to be demonstrated empirically, it is imperative to thoroughly understand various pathogenicity features of *R. pseudosolanacearum* F1C1 in host plants. Since virulence is a multifactorial phenomenon that manifests through various symptoms depending on the interaction with the host plant, using a seedling model becomes particularly useful for large-scale observation of these interactions.

## 2. Methods

### 2.1. Bacterial strains and growth conditions

The bacterial strains used in this study and their relevant characteristics are summarised in Table 1. *R. pseudosolanacearum* was cultured in BG broth (Boucher et al. 1985), composed of 0.1 % yeast extract (Hi-Media, Mumbai, India), 1 % peptone (Hi-Media), 0.1 % casamino acids (SRL, New Mumbai, India), and supplemented with 0.5 % glucose (Hi-Media) at 28 °C. *Escherichia coli* was grown in LB broth (Hi[Media) (contains 1% casein enzymic hydrolysate, 0.5% yeast extract, 1% NaCl) at 37 °C. For the preparation of solid media, 1.5 % agar (Hi-Media) was added to both BG and LB broths. The antibiotic spectinomycin (Hi-Media) was used at 50 μg/mL.

**Table 1.**
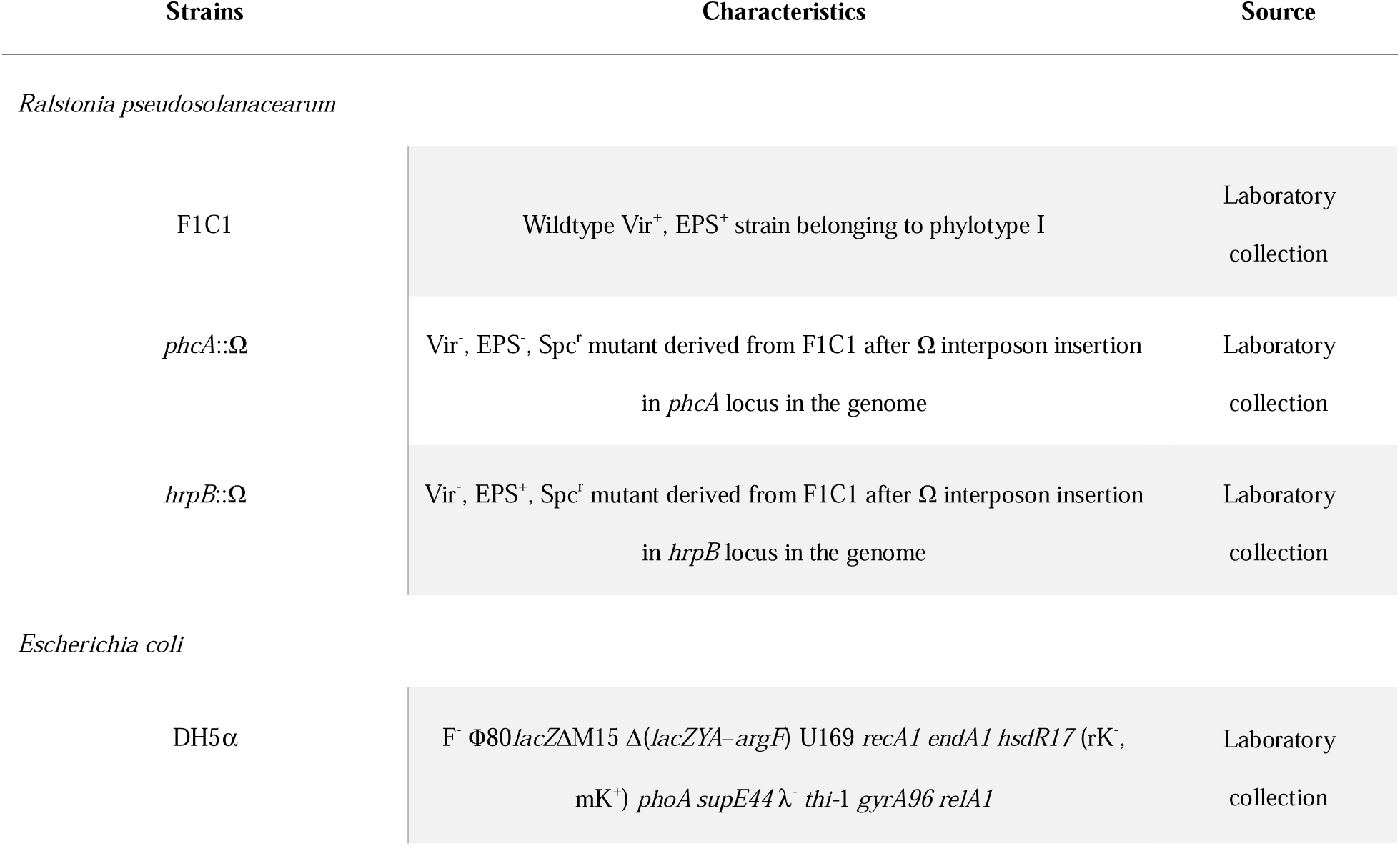
Bacterial strains used in the study.

### 2.2. Germination of seeds and growth of seedlings

Seeds of the tomato variety Pusa Ruby (Durga Seeds, U.P., India) and the eggplant variety Dev-Kiran (614) (Tokita, Karnataka, India) were employed for this study. Both tomato and eggplant seeds were germinated according to the protocols outlined by Bhuyan et al. (2025). Before sowing, tomato seeds were soaked in sterile dH_2_O at room temperature for 24 h, while eggplant seeds were soaked at 4 °C for 48 h. The seeds were subsequently sown in sterile soil.

To retain the moisture, the containers with sown seeds were tightly sealed in zip-lock plastic bags and kept in the dark at room temperature for 72 h. Following this period, the zip-lock bags were removed, and the sprouted seeds were transferred to a growth chamber (Orbitek, Scigenics, India) set at 28 °C with a 12 h photoperiod and 80 % relative humidity. The sprouted seeds were watered at regular intervals of 12 h to offset moisture loss due to transpiration and evaporation. By day 7, over 80% of the seeds from both species had germinated into two-leaf cotyledon stage seedlings. For infection assays, these 7-day-old cotyledon stage seedlings were transferred to 1.5 mL microfuge tubes containing sterile dH_2_O. Notably, prior to transfer, the roots of the seedlings were rinsed with sterile dH[O to remove residual soil adhering to them.

### 2.3. Preparation of bacterial inoculum for virulence assays

Fresh bacterial culture samples were inoculated into 10 mL of BG broth and incubated in a shaking incubator (Orbitek, Scigenics, India) maintained at 28 °C, 150 rpm for 24 h. After incubation, the bacterial suspension was centrifuged at 4000 rpm at 4 °C for 15 min (Centrifuge 5804 R, Eppendorf, Hamburg, Germany). The resulting pellet was resuspended with an equal volume of sterile dH_2_O and recentrifuged under the same conditions as above to remove any residual culture media components. The final pellet then obtained was again resuspended with an equal volume of sterile dH_2_O, and the bacterial concentration of the suspension was adjusted to approximately 10^9^ CFU/mL (OD_600_ = 1.0) (BioSpectrometer, Eppendorf, Hamburg, Germany).

### 2.4. Virulence assays in tomato and eggplant seedlings

Virulence assays were performed on 7-day-old cotyledon-stage tomato and eggplant seedlings following the methods described by Bhuyan *et al*. (2025). For leaf inoculation, the seedlings were uprooted from the soil bed and their roots were gently rinsed with sterile dH_2_O before being transferred into 1.5 mL microfuge tubes containing sterile dH_2_O. Subsequently, about one-third of the cotyledon leaves were carefully excised using sterile scissors dipped in the bacterial suspension for inoculating both leaves of the cotyledon-stage seedlings. From root inoculation, the soil-adhering roots were first cleaned and gently dried by placing the seedlings on dry tissue paper. The dried roots were then briefly dipped (for approximately 1-2 s) in the bacterial inoculum and immediately transferred to 1.5[mL microfuge tubes. After an interval of 2-5 min, sterile dH_2_O was added to each tube.

The pathogenicity assay was carried out in triplicate, with each trial comprising three sets of 40 seedlings each. In total, nine sets were used across the three experimental repetitions to assess the virulence phenotype. Seedlings were mock-inoculated with sterile dH_2_O. Additionally, seedlings inoculated with *E. coli* DH5[served as a control to assess any response of these young seedlings to a non-pathogenic Gram-negative bacterium. Pathogenicity experiments were performed at 10[CFU/mL of bacterial cell suspensions in tomato as well as eggplant seedlings. The wilting of these seedlings was recorded daily at 24-hour intervals, starting from the day of inoculation and continuing up to 10 days post-inoculation (DPI). During this 10-day observation period, the inoculated seedlings were kept in a growth chamber maintained at 28 °C with a 12 h photoperiod and 80 % relative humidity. The data presented in this study reflect the average values from three sets within a single experimental replicate. Since the remaining two replicates produced similar results, their data have not been included.

### 2.5. Statistical analysis

To evaluate the virulence of bacterial samples in host seedlings, Kaplan-Meier survival curves were plotted and log-rank tests performed using R. The R script applied in this analysis has been adapted from Bhuyan et al. (2025).

### 2.6. Isolation and confirmation of *R. pseudosolanacearum* F1C1-derived virulence-deficient mutants from wilted as well as escapee seedlings

Ten days after inoculating tomato and eggplant seedlings (10[CFU/mL) via both root and leaf methods with *phcA*::Ω and *hrpB*::Ω, randomly selected wilted and escapee seedlings from each treatment group were used for the isolation and confirmation of the inoculated bacterium, following the procedures described in Bhuyan et al. (2025), where the wild-type F1C1 strain had been previously isolated and confirmed. Seedlings were surface-sterilized, homogenized in sterile dH_2_O using sterile micropestles, and the homogenate was serially diluted up to 10[-fold, and 20 μL of the 10[-fold diluted suspension was spotted on BG-Agar medium. Early bacterial growth was assessed after 18 h, and the microcolonies were evaluated for twitching motility using an inverted phase contrast microscope. After 38 h of incubation, colony morphology was recorded, and representative isolates were subjected to multiplex PCR targeting the 16S-23S rRNA gene internal transcribed spacer region using phylotype-specific primers. DNA template preparation via alkaline lysis using NaOH, PCR reactions, and electrophoresis on 2% agarose gel with appropriate controls was performed as previously described (Bhuyan et al. 2025). Gel images were visualized and documented using a Gel-Doc system.

## 3. Results

### 3.1. Phenotypic effects of *R. pseudosolanacearum* F1C1 infection in host seedlings

To investigate the pathological dynamics of *R. pseudosolanacearum* F1C1 in practice, we conducted experiments that examined *R. pseudosolanacearum* F1C1 interaction in tomato seedlings (variety Pusa Ruby, Durga Seeds, Uttar Pradesh, India) and eggplant seedlings (variety Dev-Kiran (614), Tokita, Karnataka, India) using leaf and root inoculation methods. Upon close observation of the inoculated seedlings, apart from the usual symptom of the drooping of the stem, a plethora of phenotypes were also observed in these tomato and eggplant seedlings under study. These phenotypes include: (i) blackening of the stem, (ii) softening and translucency of the stem, (iii) emergence of lateral roots, (iv) emergence of true leaves, (v) curling of the cotyledon leaves, (vi) chlorosis of the cotyledon leaves, and (vii) blackening of the clipped cotyledon leaf tip (Fig. 1). It is noteworthy that the emergence of lateral roots occurs above the softened and translucent section of the stem. This emergence of lateral roots aids in discarding the primary root attached to the diseased portion of the stem, contributing to the plant’s effort to escape the disease. Similarly, the emergence of true leaves also aids in escaping disease by discarding the diseased cotyledon leaves. To our knowledge, these phenotypes have not been adequately documented earlier in seedlings infected by *R. pseudosolanacearum*.

**Figure 1.**
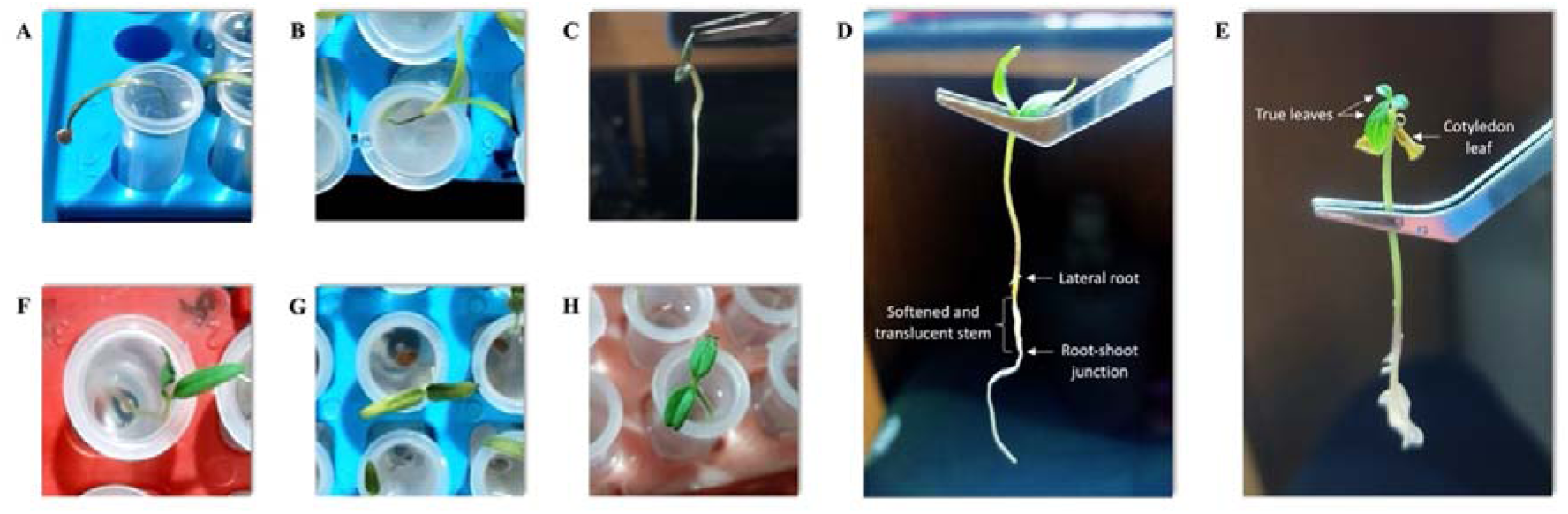
Symptoms of *R. pseudosolanacearum* F1C1 infection in seedlings of tomato and eggplant. [**A**] Drooping of the stem. [**B**] Blackening of the stem. [**C**] Softening and translucency of the stem. [**D**] Emergence of lateral roots. [**E**] Emergence of true leaves. [**F**] Curling of the cotyledon leaves. [**G**] Chlorosis of the cotyledon leaves. [**H**] Blackening of the clipped cotyledon leaf tip.

### 3.2. Morphology of softening and translucency in the stems of tomato and eggplant seedlings as a result of *R. pseudosolanacearum* F1C1 infection

Among these above phenotypes, the softening and translucency of the stem are particularly intriguing because the seedlings appeared fully intact with healthy cotyledon leaves. Therefore, this observation was unexpected. This could be observed while inspecting individual plants that appeared healthy, where the increased transparency of the stem region was noticeable (Fig. 2). Safranin staining of the seedlings was also performed for visualization of the softened and translucent portion of the stem (Supplementary Fig. 1).

**Figure 2.**
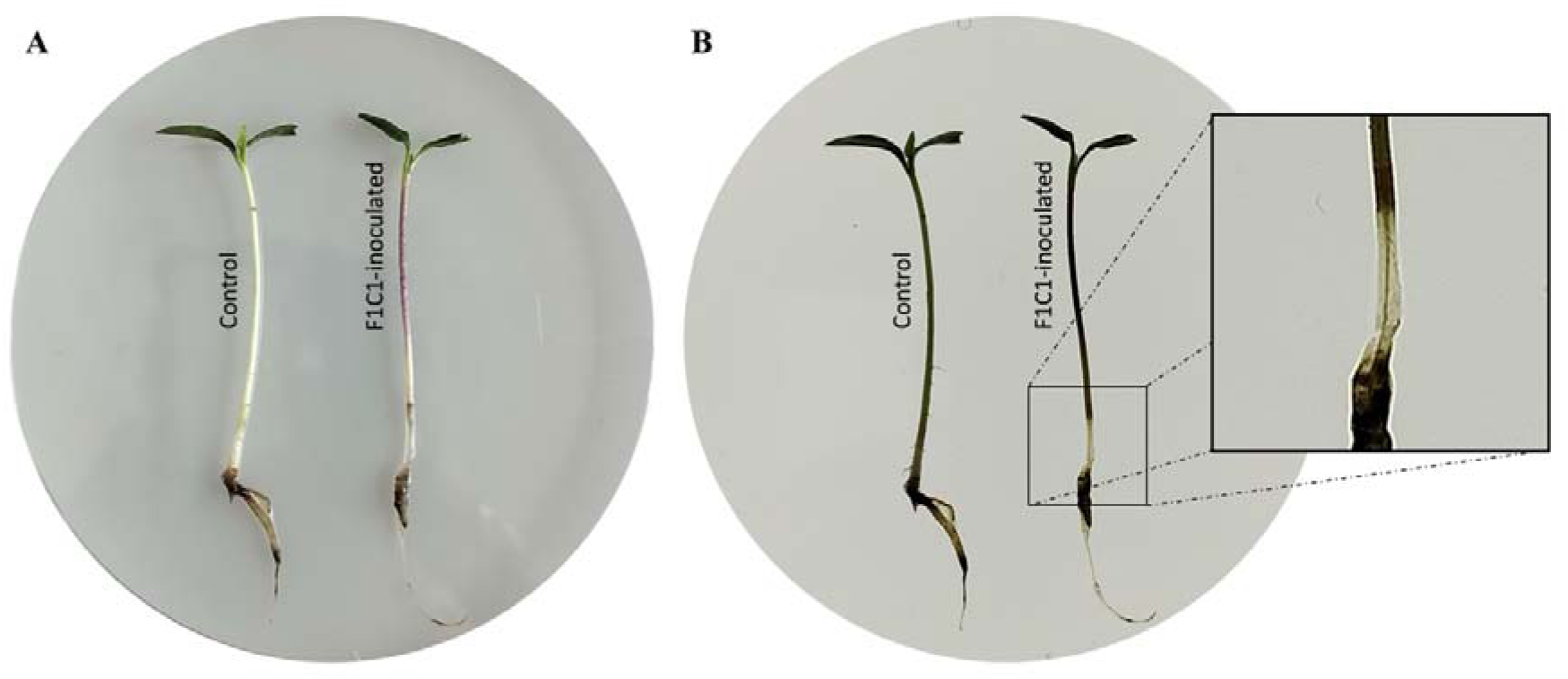
*R. pseudosolanacearum* F1C1-induced morphology of softening and translucency in seedling stems. [**A**] under normal light conditions. [**B**] on a backlit LED board exposing the intact stele of the diseased stem region.

Upon closer examination of the softened and translucent stem under 40X magnification with a compound microscope it was found that the softened and translucent seedling stems had a clear epidermal and cortex region compared to the stems of an uninfected seedling. However, the softened and translucent seedling stems appeared to maintain a healthy stele similar to the uninfected ones (Fig. 3). Notably, this phenotype was absent in all control plants that were not inoculated with *R. pseudosolanacearum* F1C1.

**Figure 3.**
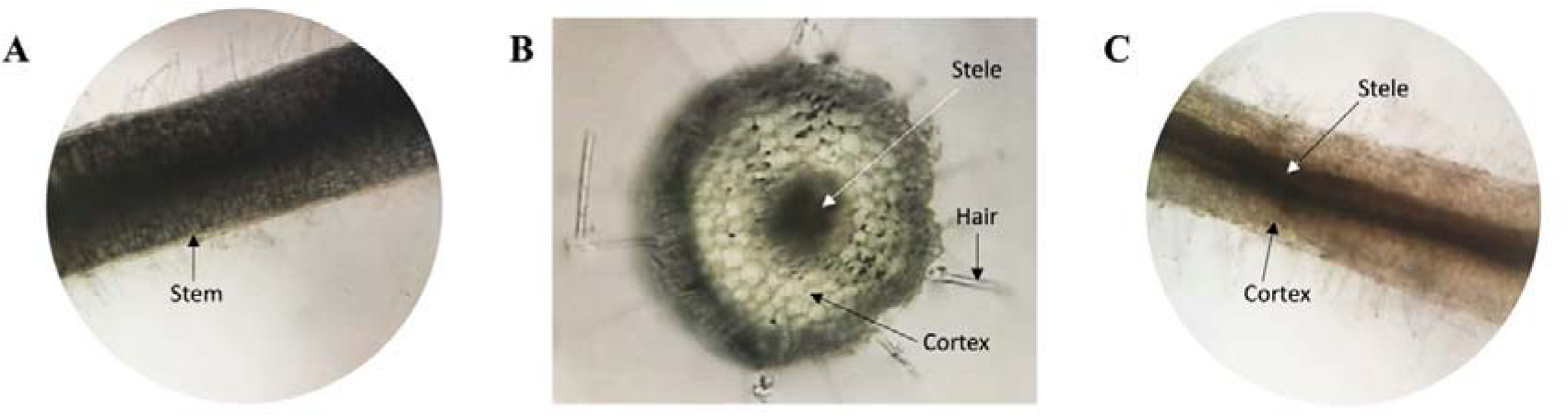
Morphology of softening and translucency of infected seedling stems observed under 40X magnification. [**A**] A healthy stem. [**B**] T.S. of a healthy stem. [**C**] An infected stem.

It is pertinent to note that the morphology of softening and translucency of the stem is predominantly observed in seedlings when *R. pseudosolanacearum* F1C1 is inoculated through both the roots and the leaves. This softened and translucent stem phenotype is restricted to the stem region that remains submerged in water, which makes it challenging to detect unless the seedling is removed and examined against a light source. The portion of the stem that is above water is visibly affected by the disease, showing signs of blackening (Fig. 4).

**Figure 4.**
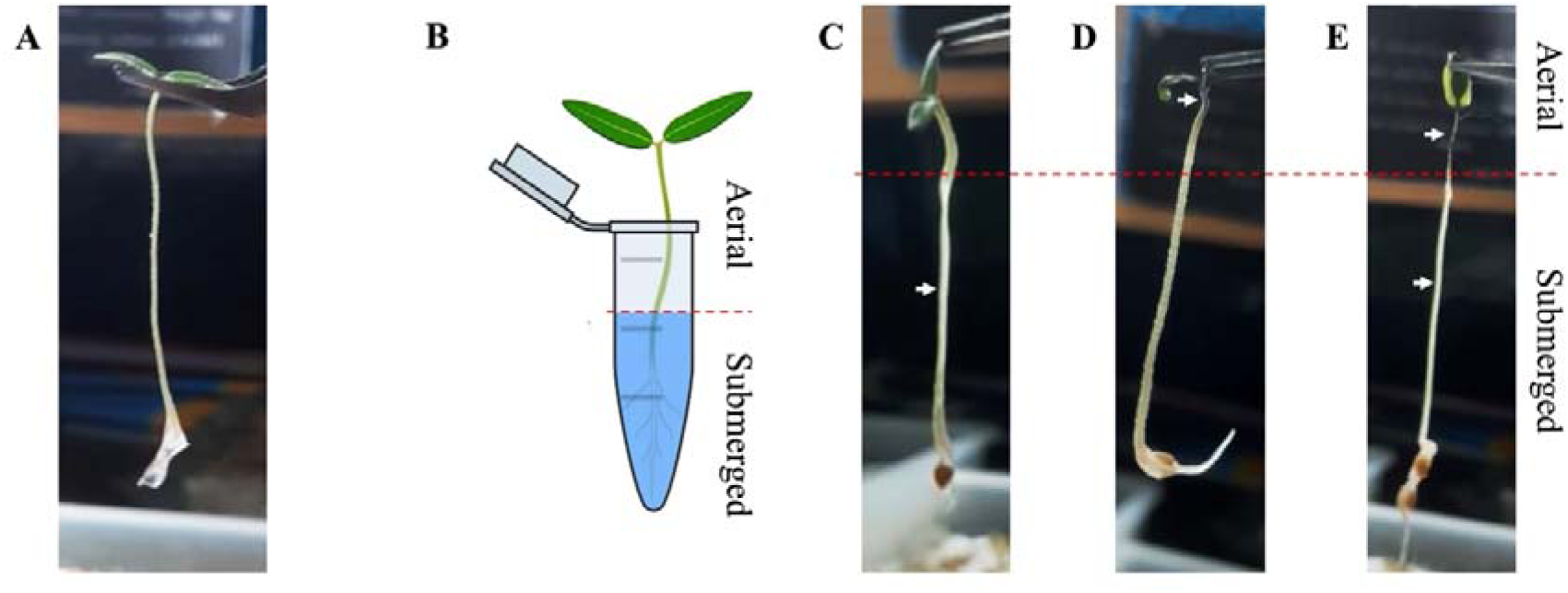
The phenotypes of stem softening and translucency, along with blackening, in *R. pseudosolanacearum* F1C1-infected seedlings. [**A**] Uninfected healthy seedling stem. [**B**] Graphical representation of the cotyledons’ aerial and submerged stem sections, illustrating where the phenotype of blackening and the phenotype of softening and translucency occur, respectively. [**C**] The morphology of softening and translucency of the submerged part of the infected seedling stem. [**D**] Blackening of the aerial part of the infected seedling stem. [**E**] The phenotypes of stem softening and translucency, along with blackening, in a single infected seedling. White arrows indicate the diseased region of the seedlings.

### 3.3. The phenotype of stem softening and translucency in virulence-deficient mutants of R. pseudosolanacearum F1C1

Tomato and eggplant seedlings were inoculated with *R. pseudosolanacearum* F1C1 and its virulence-deficient mutants, *phcA*::Ω and *hrpB*::Ω, using leaf and root inoculation methods. In the case of the wild-type F1C1, seedling death was observed by 3 DPI in tomato seedlings through both leaf and root inoculation, and in eggplant seedlings only following leaf inoculation. For *phcA*::Ω, seedling death began from 4 DPI in tomato seedlings via both inoculation routes, and in eggplant seedlings only when inoculated through the leaves (Fig. 5 & 6). In contrast, seedlings inoculated with *hrpB*::Ω did not exhibit any seedling deaths in either host through both inoculation methods. However, a slight variation in standard deviation at 6 DPI in eggplant seedlings inoculated via the leaf suggests the incidence of an isolated seedling death, though the average remained zero (Fig. 6). These observations reaffirm the high susceptibility of eggplant seedlings when inoculated through the leaves, as discussed by us earlier (Bhuyan et al. 2025).

**Figure 5.**
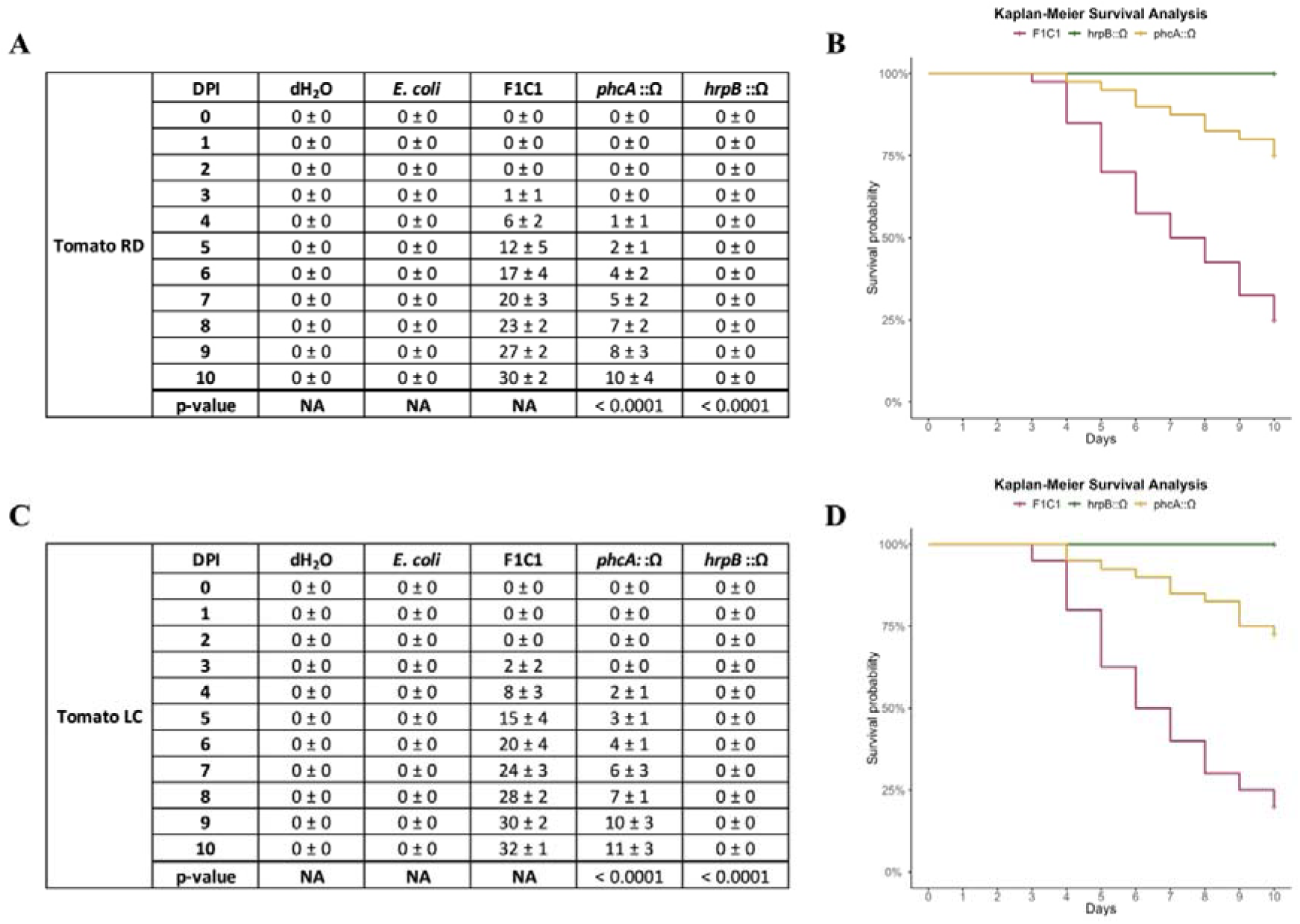
Virulence assessment in tomato seedlings. [**A** & **C**] Seedling deaths following inoculation of bacterial samples through root and leaf, respectively. [**B** & **D**] Kaplan-Meier survival analysis of tomato seedlings inoculated with bacterial samples via root and leaf.

**Figure 6.**
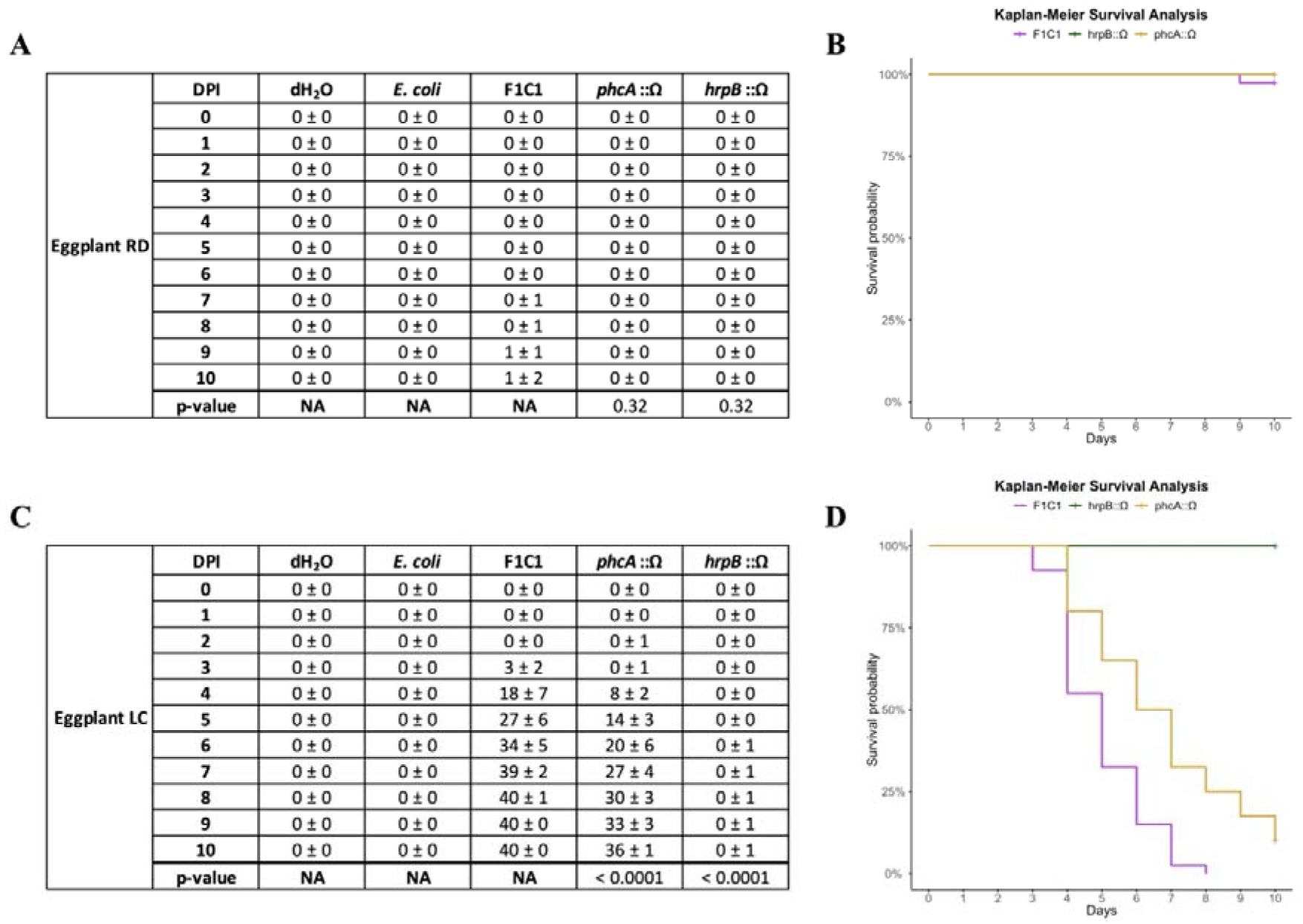
Virulence assessment in eggplant seedlings. [**A** & **C**] Seedling deaths following inoculation of bacterial samples through root and leaf, respectively. [**B** & **D**] Kaplan-Meier survival analysis of eggplant seedlings inoculated with bacterial samples via root and leaf.

The *hrpB*::Ω mutant was essentially avirulent, while *phcA*::Ω displayed weak virulence in tomato but caused a higher rate of seedling death in eggplant when inoculated through the leaf (Fig. 5 & 6). This is consistent with the findings of Phukan et al. (2019). The virulence of the wild-type F1C1, as observed in tomato seedlings through both leaf and root inoculation and in eggplant seedlings via the leaf, was consistent with our earlier findings (Bhuyan et al. 2025). As previously noted, eggplant seedlings were notably less susceptible to all tested bacterial strains when inoculated via the roots.

Observations during the virulence assays indicated that the onset of the phenotype of stem softening and translucency typically began at 5 DPI, regardless of the inoculation route, though it was also occasionally observed as early as 4 DPI. Notably, this was frequently observed in both tomato and eggplant seedlings inoculated with the wild-type F1C1, with eggplant showing a notably higher incidence overall (≅ 60 %) across both root and leaf inoculation methods than tomato (≅ 20 %). Seedlings inoculated with the *phcA*::Ω mutant exhibited a reduced occurrence of this phenotype (≅ 10 %) in both tomato and eggplant when inoculated through the roots, and in tomato following inoculation through leaves. However, the phenotype became substantially more evident in eggplant following leaf inoculation, where a noticeably higher proportion of seedlings (≅ 50 %) displayed the symptom. Interestingly, the *hrpB*::Ω mutant, while not inducing any visible symptoms in tomato, was able to elicit occasional instances (≅ 10 %) of softening and translucency in eggplant seedlings (Fig. 7). These findings further underscore the higher susceptibility of eggplant seedlings as a host, highlighting that even the virulence-deficient *hrpB*::Ω mutant, which fails to produce visible symptoms in tomato, was capable of inducing disease-associated phenotypes in eggplant.

**Figure 7.**
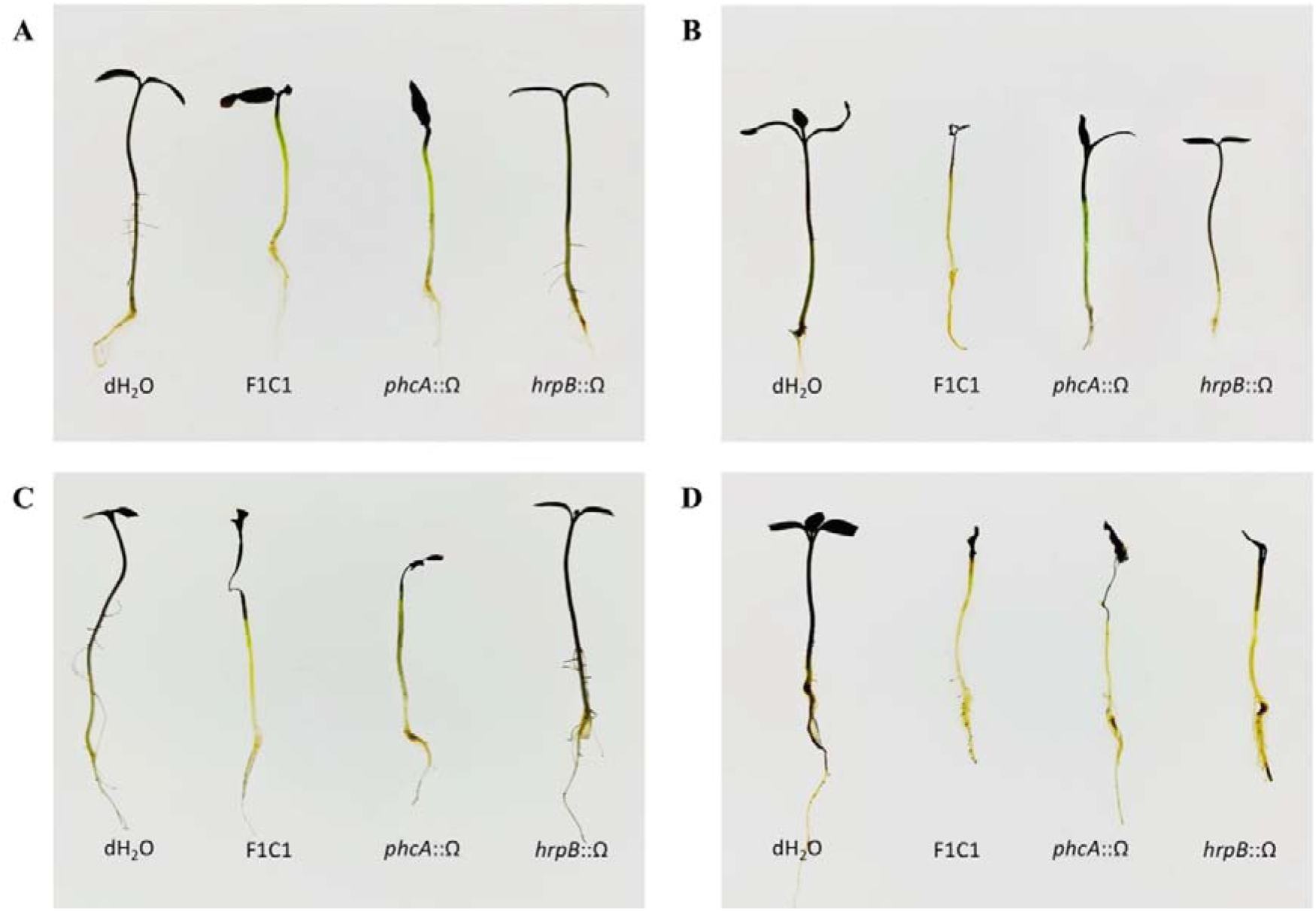
Symptom of stem softening and translucency upon inoculation with bacterial samples. [**A** & **C**] Tomato seedlings through root and leaf, respectively. [**B** & **D**] Eggplant seedlings through root and leaf, respectively.

### 3.4. Confirmation of the presence of *R. pseudosolanacearum* F1C1-derived virulence-deficient mutants in both wilted as well as escapee seedlings of tomato and eggplant

The presence of the phytopathogen in both wilted and escapee tomato seedlings inoculated through the leaf was demonstrated using the *mCherry*-tagged F1C1 through fluorescence microscopy as well as by the isolation of the bacterium from the seedlings (Phukan 2018). The *GUS*-tagged F1C1 was also used to further confirm the presence of phytopathogen in the seedlings (Phukan 2018). We performed additional experiments to study the presence of *R. pseudosolanacearum* F1C1 in root-inoculated wilted and escapee seedlings of tomato and eggplant, as well as leaf-inoculated wilted and escapee tomato seedlings. No escapees were observed in the case of leaf-inoculated eggplant seedlings due to their high susceptibility, and therefore, *R. pseudosolanacearum* F1C1 could only be isolated from wilted eggplant seedlings (Bhuyan et al. 2025). In this manuscript, we only confirm the presence of *R. pseudosolanacearum* F1C1-derived virulence-deficient mutants of *phcA*::Ω and *hrpB*::Ω in the inoculated seedlings of both hosts.

On 10 DPI, the wilted and escapee seedlings of tomato and eggplant were homogenized, and 20 μL of 10^5^-fold serially diluted homogenate was spotted on BG-Agar plates supplemented with glucose and incubated at 28 °C to facilitate bacterial growth. After 18 h of incubation, the spotted area was examined for the presence of microcolonies. These sparsely scattered microcolonies exhibited twitching motility characteristic of the early growth phase of *R. pseudosolanacearum,* forming circular, mesh-like structures (Fig. 8), unlike elongated microcolonies that are typically observed at higher bacterial concentrations (Bhuyan et al. 2024). By 38 h of incubation, distinct colony morphologies were observed on the BG-Agar plates: the *hrpB*::Ω mutant formed white, opaque, fluidal colonies, whereas the *phcA*::Ω mutant produced white, translucent, afluidal colonies. Both types were surrounded by thin, translucent, concentric rings indicative of active bacterial twitching motility, and were visible to the naked eye. These colony morphologies were characteristic of *R. pseudosolanacearum*, suggesting the successful presence and proliferation of the inoculated strain. Further assessment of the representative *R. pseudosolanacearum* F1C1-derived virulence-deficient mutant colonies using phylotype-specific multiplex PCR, followed by agarose gel electrophoresis, revealed amplified DNA bands at 144 bp. This result confirmed that the colonies isolated from both wilted and escapee seedlings of tomato and eggplant belong to phylotype I of the RSSC, indicating that they were indeed the virulence-deficient mutants derived from *R. pseudosolanacearum* F1C1 (Fig. 8).

**Figure 8.**
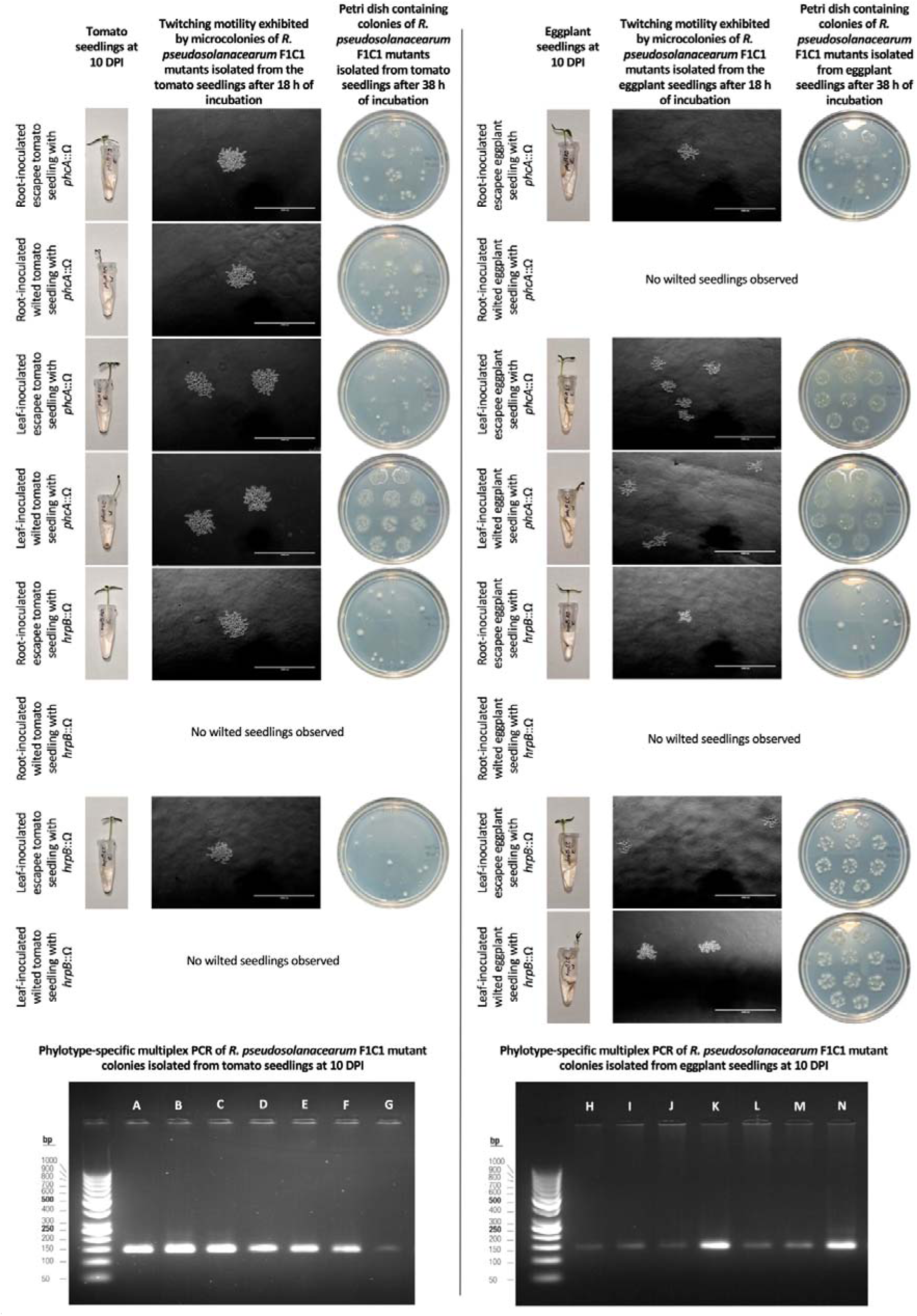
Isolation and confirmation of *R. pseudosolanacearum* F1C1 from both wilted and escapee seedlings of tomato and eggplant on 10 DPI through seedling homogenization, dilution spotting and subsequent assessment of twitching motility of microcolonies, morphology of grown-up colonies and phylotype-specific multiplex PCR targeting the 16S– 23S rRNA region. Lane A & H – *R. pseudosolanacearum* F1C1 (lab collection), Lane B – bacteria isolated from an escapee tomato seedling root-inoculated with *phcA*::Ω, Lane C – bacteria isolated from a wilted tomato seedling root-inoculated with *phcA*::Ω, Lane D – bacteria isolated from an escapee tomato seedling leaf-inoculated with *phcA*::Ω, Lane E – bacteria isolated from a wilted tomato seedling leaf-inoculated with *phcA*::Ω, Lane F – bacteria isolated from an escapee tomato seedling root-inoculated with *hrpB*::Ω, Lane G – bacteria isolated from an escapee tomato seedling leaf-inoculated with *hrpB*::Ω, Lane I – bacteria isolated from an escapee eggplant seedling root-inoculated with *phcA*::Ω, Lane J – bacteria isolated from an escapee eggplant seedling leaf-inoculated with *phcA*::Ω, Lane K – bacteria isolated from a wilted eggplant seedling leaf-inoculated with *phcA*::Ω, Lane L – bacteria isolated from an escapee eggplant seedling root-inoculated with *hrpB*::Ω, Lane M – bacteria isolated from an escapee eggplant seedling leaf-inoculated with *hrpB*::Ω, Lane N – bacteria isolated from a wilted eggplant seedling leaf-inoculated with *hrpB*::Ω.

## 4. Discussion

*R. pseudosolanacearum* carries a multitude of virulence determinants such as flagella-dependent swimming motility (Tans-kersten et al. 2004); type 4 pili-mediated twitching motility (Bhuyan et al. 2023; Bhuyan et al. 2024); type II protein secretion system that secretes a consortium of cell wall degrading enzymes such as polygalacturonases, pectin methyl esterases, endoglucanases and cellobiohydrolases (Liu et al. 2005); exopolysaccharides (Hayashi et al. 2019a); adhesins such as lectins and filamentous hemagglutinins (Kumar 2014; Hayashi et al. 2019b; Kostlanova et al. 2005); type III secretion system that secretes various type 3 effector proteins to overcome the host plant’s defense mechanism (Valls et al. 2006); efflux pumps to efflux out toxic substances (Brown et al. 2007); etc. The expression levels of these virulence factors can vary, potentially leading to different pathological phenotypes in the host. Understanding these pathological phenotypes and the different modes of pathogenesis is crucial, as virulence is a multifactorial phenomenon with various factors contributing to pathogenicity in different degrees depending on their expression and function within the host plant. Therefore, associating these phenotypes with the magnitude of virulence function expression is essential for unravelling the complexity of *R. pseudosolanacearum* F1C1 infection.

The observation of stem softening and translucency as a pathological feature restricted to the water-submerged regions of cotyledon-stage seedlings represents a novel phenotype not only in tomato and eggplant but also in chilli (data not presented). This is because the softened tissues in the submerged regions are supported by the buoyancy of water, allowing them to retain their shape and appear translucent, whereas similar tissues above water tend to collapse, blacken, and dry before the translucency can be observed. Microscopic analyses of the softened, translucent stem region further enrich our understanding. The cortex and epidermal layers of these stems appeared notably more translucent compared to control tissues, while the stele remained largely intact and structurally preserved. The morphological changes induced by this bacterium in seedlings during the infection process have been reported earlier limited to the root region only. Vasse et al (1995) reported that the bacterium colonizes the epidermis and readily multiplies in the intercellular spaces of the cortex. In agreement with this, Inoue et al. (2023) demonstrated that *R. pseudosolanacearum* invades and colonises the intercellular spaces of the epidermis and cortex of the plant tissues, wherein the infectivity is aided by plant cell wall-degrading enzymes secreted by the type II protein secretion system. Further, Digonnet et al. (2012) reported a rapid plasmolysis of the epidermal, cortical, and endodermal cells, further supporting our observations. It is pertinent to note that in all these studies, the damage rendered to the root regions by the pathogen could only be microscopically observed after cross-sectioning the root regions. But the current study demonstrates the damage in the internal region of the stem portion noticed externally with the unaided eye. Taken together, these reports, along with our current results of selective stem degradation in seedlings, suggest that *R. pseudosolanacearum* F1C1 is capable of discriminating among plant tissues, possibly led by specific enzyme activity or intrinsic tissue properties. Such tissue-selective disintegration may allow the pathogen to optimize its spread and nutrient acquisition while preserving the vascular integrity necessary for systemic colonization.

Consistent with the pattern observed for the wild-type F1C1 strain, the capacity of the *hrpB*::Ω mutant to induce this novel phenotype in ≅ 10 % of eggplant but not in tomato further emphasizes the differential susceptibility between the two hosts. Despite being impaired in its type III secretion system and thus avirulent, *hrpB*::Ω nonetheless triggers cortex disintegration and visible stem softening and translucency in eggplant. This again points to the higher vulnerability of eggplant seedlings towards *R. pseudosolanacearum* F1C1 infection. The *hrpB*::Ω mutant presents yet another interesting observation. Although it retains exopolysaccharide production *in vitro* (data not presented), it is avirulent in either host. Nevertheless, its persistence within both tomato and eggplant seedlings at 10 DPI, especially in the escapee seedlings, demonstrates its ability to survive and possibly even colonize internal tissues. To this day, no published study has directly quantified exopolysaccharide levels *in planta* for *hrpB* mutants of RSSC. This, along with future comparative assessment of its *in planta* growth rate in susceptible hosts, would also help us gain clarity in this regard, as exopolysaccharide secretion is a density-dependent phenomenon.

Conversely, the *phcA*::Ω mutant, deficient in exopolysaccharide production yet capable of causing significant seedling death in eggplant via leaf inoculation, poses critical questions regarding the canonical role of exopolysaccharides in virulence. Exopolysaccharides have long been regarded as a central virulence factor in RSSC, contributing to vascular occlusion and immune evasion. However, our findings suggest that even in the absence of exopolysaccharides, high seedling death is possible, particularly in susceptible hosts. The shared symptomatology of seedling stem softening and translucency between infections caused by the wild-type F1C1 strain and the exopolysaccharide-deficient *phcA*::Ω mutant, due to cortex disintegration, strongly implicates other virulence components, such as cell wall-degrading enzymes, to play a central role in *R. pseudosolanacearum-* induced seedling death.

## Ethics

Our work does not contain experiments involving animals and/or human participants.

## Data accessibility

The article has no additional data.

## Declaration of AI use

We have not used AI-assisted technologies in creating this article.

## Authors’ contributions

S.B.: Investigation, methodology, formal analysis, software, validation, visualisation, writing-original draft, writing-review and editing; L.B./M.J.: Investigation, methodology, visualisation, validation, writing-review and editing; S.Begum/S.J.G./L.D.: Investigation, validation, writing-review and editing; S.K./S.C.: Investigation, validation; M.M.: Supervision, writing-review and editing; S.K.R.: Conceptualization, supervision, visualisation, writing-original draft, writing-review and editing.

## Acknowledgements

S.B. is thankful for the JRF/SRF fellowship from the UGC-NFSC, GoI, New Delhi.

M.J. is thankful to Tezpur University for the institutional fellowship followed by the JRF fellowship from CSIR, GoI, New Delhi. L.B. and S.C. are thankful to DBT, GoI for the MSc fellowship and project research grant. S.Begum and S.J.G. are thankful for the JRF fellowship from the DBT, GoI, New Delhi grant (BT/PR41637/NER/95/1753/2021) awarded to S.K.R. L.D. is thankful to DST, GoI, New Delhi for the INSPIRE-JRF fellowship. S.K. is thankful for the JRF fellowship from UGC, GoI, New Delhi.

