## Supplementary data for "Morphology of softening and translucency in the stems of tomato and eggplant seedlings due to infection by *Ralstonia pseudosolanacearum* F1C1"

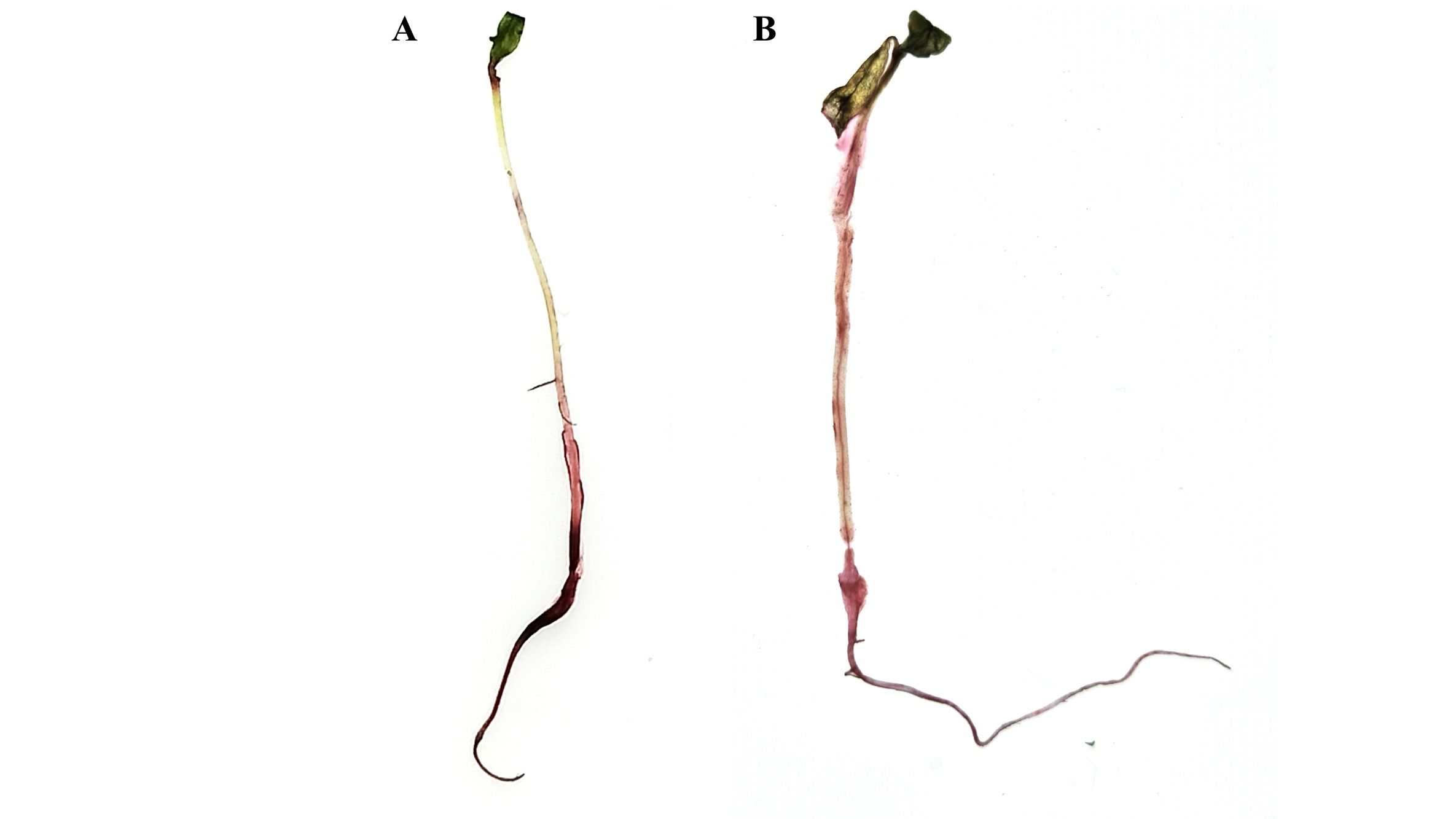

**Supplementary Figure 1.** Safranin staining of [**A**] Control seedling mock-inoculated with sterile dH_2_O, and [**B**] *Ralstonia pseudosolanacearum* F1C1-inoculated seedling to aid visualization of the softened and translucent portion of the seedling. A staining solution was prepared by diluting 250 μL of Safranin with 750 μL of dH_2_O in a 1.5 mL microcentrifuge tube. The 6-day post-inoculated test seedlings were dipped in the working staining solution for 30-40 min, until adequate coloration was observed. Excess stain was removed by rinsing the stained seedling with absolute ethanol.
